# torch-projectors: A High-Performance Differentiable Projection Library for PyTorch

**DOI:** 10.64898/2026.03.10.710885

**Authors:** Dimitry Tegunov

## Abstract

Fourier-space projection operations are central to electron microscopy single-particle analysis and electron tomography algorithms. Machine learning methods require differentiable implementations for end-to-end model training, but PyTorch’s built-in operations are too slow for practical use. This paper introduces `torch-projectors`: a high-performance library for differentiable Fourier-space projections in PyTorch. The library provides 2D and 3D forward and backward projection operators with linear and cubic interpolation, supporting gradient calculation for all inputs. Optimized for CPU, Apple Silicon (MPS), and CUDA devices, `torch-projectors` outperforms `torch-fourier-slice` by 1–2 orders of magnitude.

## 1 Introduction

Transmission electron microscopes capture 2D projections of the sample’s Coulomb potential at very high optical resolution. Multiple projections from different angles enable computational reconstruction of the original 3D potential, allowing atomic model building in favorable cases. Cryogenic electron microscopy (cryo-EM) has emerged as the dominant method for solving protein structures and investigating their dynamics. Under the thin-object approximation, the Fourier slice theorem [1] establishes that each 2D projection corresponds to a central slice through the 3D Fourier transform of the potential. This relationship underpins algorithms in electron microscopy (EM) single-particle analysis (SPA) and electron tomography (ET).

Modern machine learning methods can quickly optimize vast parameters using gradient back-propagation in differentiable models. PyTorch [2] provides automatic differentiation for implementing such models. However, many EM algorithms require differentiable Fourier-space projection operations not natively supported by PyTorch. While implementing these operations using PyTorch’s built-in functions is possible [3], performance is often insufficient and memory overhead too high for practical use. `torch-projectors` implements each projection operation in a single call without storing intermediate values, limiting memory footprint to input and output tensors.

RELION [4] popularized an accurate, computationally efficient interpolation method for Fourier-space projection operations in SPA. Forward projections sample from oversampled references (zero-padded in real space), while backward projections insert data into similarly oversampled reconstructions before cropping to regular size. This improves linear interpolation accuracy at the cost of increased memory usage. While effective for conventional SPA algorithms, machine learning methods may be memory-constrained or require costly Fourier transforms at every pass for oversampling. `torch-projectors` implements Catmull-Rom cubic interpolation [5] alongside linear interpolation, significantly improving accuracy without oversampling.

The use of dual volumes for reconstructions is another elegant approach implemented in RELION: one volume with complex-valued data and another tracking real-valued weights for each Fourier component. This enables efficient normalization of sampling counts and Wiener-like deconvolution when weights incorporate the contrast transfer function (CTF) of back-projected images. `torch-projectors` implements this approach with optional gradient back-propagation through weight components.

While 3D →2D forward and 2D →3D backward projections dominate SPA, procedures like 2D particle classification require same-dimensionality projections. `torch-projectors` implements all four operations.

The terms ‘forward’ and ‘backward’ carry dual meanings throughout the text: When referring to the direction of the projection, a forward projection samples data from a reference tensor, while a backward projection scatters data into a reconstruction tensor. In PyTorch’s automatic differentiation framework, the forward pass computes outputs from inputs, while the backward pass computes input gradients from output gradients.

## 2 Methods

### 2.1 Data Conventions

`torch-projectors` operates exclusively in Fourier space using PyTorch’s RFFT format, following FFTW conventions [6]. Unlike RELION’s implementation, tensors remain RFFT-formatted throughout projection operations without intermediate format changes.

Reconstruction and projection data must be shifted (fftshift) in real space to center at the 0th tensor element before fast Fourier transform (FFT). Results must be inverse-shifted (ifftshift) in real space to restore conventional centering.

All spatial dimensions must be square (2D) or cubic (3D) with even sizes. 2D reconstructions use shape [*B, N, N/*2 + 1] where *B* is the batch dimension. 3D reconstructions use shape [*B, D, H, W/*2 + 1]. Projections follow the same RFFT convention with shape [*B, P, N, N/*2 + 1], where *B* is the reconstruction batch dimension, and *P* is the pose dimension.

Rotations are specified as matrices: [*B, P*, 2, 2] for 2D and [*B, P*, 3, 3] for 3D. Translation shifts use shape [*B, P*, 2] and are applied via phase modulation. Positive shift values move image contents along the positive axis direction. In back-projection, shifts are applied in the opposite direction to ensure inverse relationship with forward projection. Batch broadcasting is supported: *B* can be 1 or match the reconstruction batch size, and the *P* dimension in either rotations or shifts (but not both) can be 1 for shared values across poses.

### 2.2 Interpolation

`torch-projectors` supports linear and cubic interpolation methods for sampling Fourier-space data at non-integer coordinates. Linear interpolation uses standard multilinear kernels: bilinear for 2D operations (4-point support) and trilinear for 3D operations (8-point support).

Cubic interpolation employs separable Catmull-Rom kernels with parameter *a* = −0.5, providing *C*^1^ continuity and exact interpolation through control points. The kernel function is defined as:

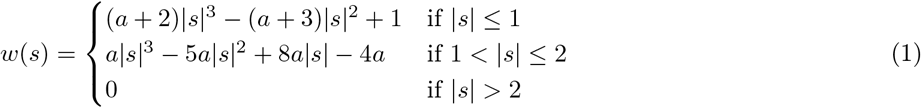

For bicubic interpolation, a 4×4 neighborhood is sampled around each coordinate, while tricubic interpolation samples a 4×4×4 neighborhood. The final interpolated value is computed as the separable product of 1D kernel evaluations along each dimension.

All interpolation operations support oversampling, where coordinates are scaled by factors *>* 1 to sample from reconstructions previously zero-padded in real space. This improves interpolation accuracy at constant computational load but increases memory usage. The sampling grid remains sparse, matching the unpadded reconstruction box size, eliminating the need for additional real-space cropping.

Optional hard low-pass filtering can be applied during projection operations, and the projection dimensions can be made smaller than the reference to downsample the resulting real-space image.

Shifts are applied through phase modulation of complex Fourier components according to the Fourier shift theorem: *F* {*f* (**x** − **s**)*}* = *F {f* (**x**)} · *e*^−*i*2*π***k**·**s**^, where **k** are Fourier coordinates and **s** is the shift vector.

### 2.3 Friedel Symmetry

The Fourier transform of a real-valued function exhibits Friedel symmetry, characterized by the relationship:

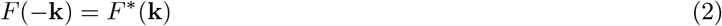

where *F* ^∗^(**k**) denotes the complex conjugate at frequency **k**. This symmetry property allows PyTorch’s RFFT format to store only the non-negative frequency components along the last dimension, yielding the characteristic […, *N/*2 + 1] shape.

However, the zero-frequency axis (where *k*_*c*_ = 0) requires careful handling during projection operations to maintain mathematical consistency and prevent double-counting:

#### Forward Pass

Standard Friedel symmetry lookup is applied when sampling negative *k*_*c*_ coordinates. For *k*_*c*_ *<* 0, the implementation accesses *F* (−*k*_*c*_, −*k*_*r*_) and returns its complex conjugate *F* ^∗^(−*k*_*c*_, −*k*_*r*_) according to equation 2.

#### Backward Pass

To avoid over-representation in gradient accumulation, the implementation skips processing the negative half of the zero-frequency axis (specifically, components where *k*_*c*_ = 0 and *k*_*r*_ *<* 0), since these are present twice in the RFFT-formatted gradient.

#### Accumulation on Zero-Frequency Axis

When projection operations result in values that must be accumulated at positions on the zero-frequency axis, the implementation maintains Friedel symmetry by inserting contributions at both the target location (*k*_*r*_, 0) and its symmetric counterpart (−*k*_*r*_, 0).

### 2.4 Implementation Architecture

`torch-projectors` follows PyTorch’s hybrid C++/Python extension architecture using the `TORCH_LIBRARY` registration system. The library implements a multi-backend design with separate optimized kernels for CPU, Apple Silicon (MPS), and CUDA devices.

The Python interface wraps C++ operators with custom `torch.autograd.Function` classes that handle gradient computation. Library initialization loads the compiled C++ extension and registers Python autograd functions using `torch.library.register_autograd`. This architecture provides seamless integration with PyTorch’s automatic differentiation while maintaining high performance through backend-specific optimizations.

### 2.5 2D → 2D Forward Projection

The 2D forward projection operator generates rotated 2D projections from 2D Fourier-space reconstructions. Given a reconstruction tensor [*B, N, N/*2+1] and rotation matrices [*B*_*rot*_, *P*, 2, 2], the operator produces projections [*B, P, N*_*out*_, *N*_*out*_*/*2+ 1] by sampling the reconstruction at transformed Fourier coordinates.

#### 2.5.1 Forward Pass

For each output Fourier coordinate **k**_*out*_ in the projection, the operator computes the corresponding coordinate in the reconstruction as:

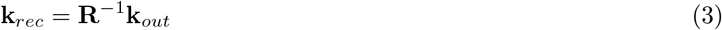

where **R** is the 2D rotation matrix. The projection value is obtained by interpolating the reconstruction at **k**_*rec*_, then applying phase modulation for translation:

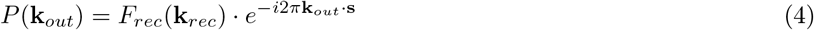

where **s** is the shift vector.

#### 2.5.2 Backward Pass

During backpropagation, the incoming projection gradients are first phase-modulated using the conjugate shift:

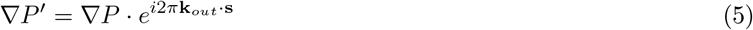

Reconstruction gradients are then accumulated via transposed interpolation operations at the transformed coordinates **k**_*rec*_.

Analytical rotation matrix gradients require careful application of the chain rule: Since **k**_*rec*_ = **R**^−1^**k**_*out*_, the gradient with respect to rotation matrix element *R*_*ij*_ is:

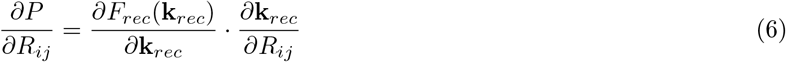

For bilinear interpolation, spatial derivatives are computed from the 2×2 sample grid using separable linear kernel derivatives:

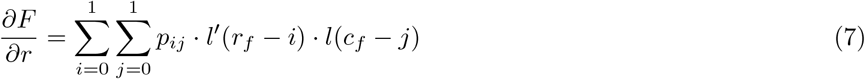

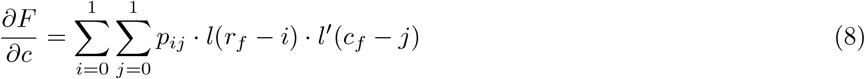

where *l*(*s*) = 1 − |*s*| for |*s*| ≤ 1 and *l*^*′*^(*s*) = sign(*s*) for |*s*| *<* 1.

For bicubic interpolation, derivatives follow from separable Catmull-Rom kernel derivatives over the 4×4 neighborhood:

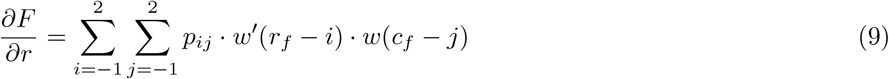

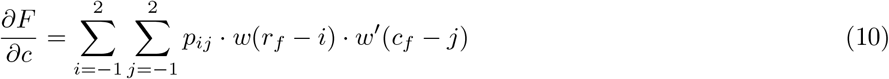

where *w*^*′*^(*s*) is the cubic kernel derivative. The coordinate transformation derivatives 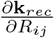 are computed from the matrix inverse relationships.

Shift gradients follow from the phase modulation derivative:

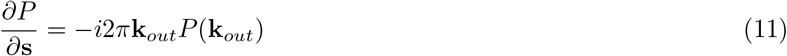

### 2.6 2D → 2D Backward Projection

The 2D backward projection operator is the mathematical adjoint of the forward projection operation. It accumulates 2D Fourier-space projections [*B, P, N, N/*2 + 1] into 2D reconstructions [*B, N*_*rec*_, *N*_*rec*_*/*2 + 1], where the reconstruction size accounts for oversampling: *N*_*rec*_ = *N* · oversampling.

#### 2.6.1 Forward Pass

For each projection coordinate **k**_*proj*_ = (*k*_*r*_, *k*_*c*_), the corresponding coordinate in the 2D reconstruction is computed using equation 3 with the same 2×2 rotation matrix **R**. The projection value is then accumulated into the reconstruction at the transformed coordinate. When shifts are present, the projection data is first conjugate phase-modulated:

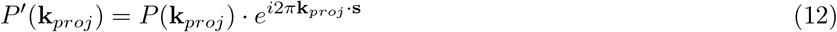

where the conjugate phase ensures the mathematical adjoint relationship with forward projection.

The accumulation process uses 2D interpolation kernels in transpose mode. For bilinear interpolation, contributions are distributed to the 2×2 neighborhood around each fractional coordinate. For bicubic interpolation, contributions are spread across the 4×4 neighborhood according to the Catmull-Rom weights defined in equations 7–8 and equations 9.

Optional weight accumulation supports applications requiring per-Fourier component weights, such as CTF correction in cryo-EM. Weights are accumulated using the absolute values of the 2D interpolation kernel weights, maintaining a reference for downstream normalization (e.g. for Wiener-like filters).

#### 2.6.2 Backward Pass

During backpropagation, projection gradients are computed using forward projection of reconstruction gradients, exploiting the adjoint relationship. Rotation matrix gradients follow equation 6 with spatial derivatives computed using equations 7–8 for bilinear interpolation or equations 9 for bicubic interpolation. Shift gradients are computed using equation 11, applied to the accumulated reconstruction data.

### 2.7 3D → 2D Forward Projection

The 3D→ 2D forward projection operator implements the Central Slice Theorem, generating 2D projections from 3D Fourier-space reconstructions. Given a 3D reconstruction tensor [*B, D, H, W/*2 + 1], 3×3 rotation matrices [*B, P*, 3, 3], and optional 2D shifts [*B, P*, 2], the operator produces projections [*B, P, H*_*out*_, *W*_*out*_*/*2 + 1] by sampling central slices through the 3D volume.

#### 2.7.1 Forward Pass

For each output projection coordinate **k**_*proj*_ = (*k*_*r*_, *k*_*c*_), the operator extends it to 3D by setting the third coordinate to zero: **k**_3*D*_ = (*k*_*c*_, *k*_*r*_, 0). This implements the central slice through the origin required by the Central Slice Theorem.

The corresponding coordinate in the 3D reconstruction is computed using the 3×3 rotation matrix:

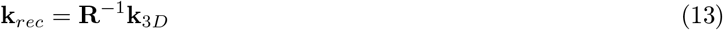

where **R** is the 3×3 rotation matrix. The projection value is obtained by trilinear or tricubic interpolation in the 3D volume at **k**_*rec*_, then applying phase modulation for translation:

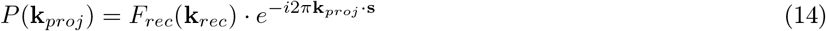

where **s** is the 2D shift vector applied to the projection coordinates.

#### 2.7.2 Backward Pass

During backpropagation, the incoming projection gradients are first phase-modulated using the conjugate shift applied to the 2D projection coordinates (using equation 5).

Reconstruction gradients are then accumulated via 3D transposed interpolation operations at the transformed coordinates **k**_*rec*_.

Analytical 3×3 rotation matrix gradients require extension of the chain rule to three dimensions: Since **k**_*rec*_ = **R**^−1^**k**_3*D*_, the gradient with respect to rotation matrix element *R*_*ij*_ is:

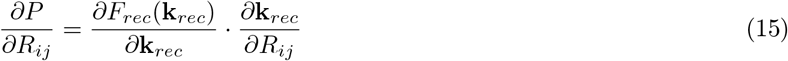

For trilinear interpolation, spatial derivatives are computed from the 2×2×2 sample grid using separable linear kernel derivatives:

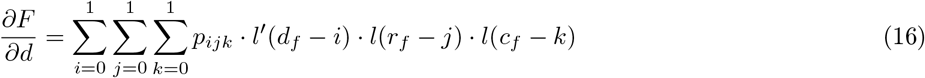

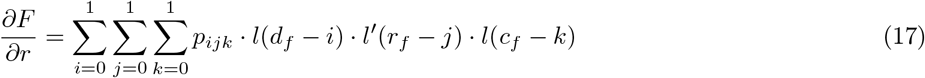

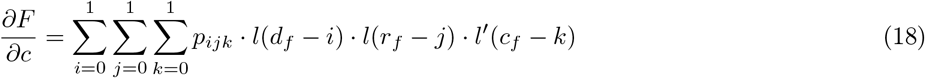

where *l*(*s*) = 1 − |*s*| for |*s*| ≤ 1 and *l*^*′*^(*s*) = sign(*s*) for |*s*| *<* 1.

For tricubic interpolation, derivatives follow from separable Catmull-Rom kernel derivatives over the 4×4×4 neigh-borhood:

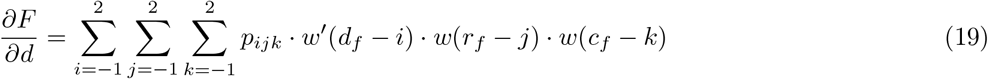

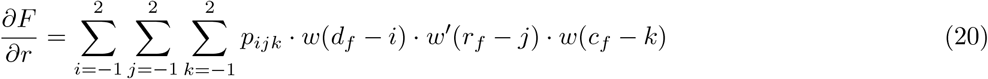

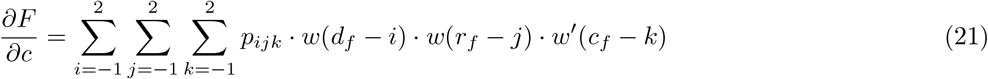

where *w*^*′*^(*s*) is the cubic kernel derivative. The coordinate transformation derivatives 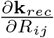 are computed from the matrix inverse relationships.

Shift gradients follow from the phase modulation derivative applied to the 2D projection coordinates:

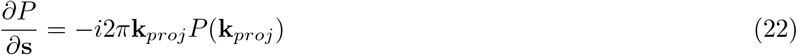

### 2.8 2D → 3D Backward Projection

The 2D →3D backward projection operator is the mathematical adjoint of the 3D →2D forward projection operation. It accumulates 2D Fourier-space projections [*B, P, H, W/*2+1] into 3D reconstructions [*B, D, H*_*rec*_, *W*_*rec*_*/*2+1], where the reconstruction dimensions account for oversampling and form a cubic volume: *D* = *H*_*rec*_ = *W*_*rec*_ = *H* · oversampling.

#### 2.8.1 Forward Pass

For each projection coordinate **k**_*proj*_ = (*k*_*r*_, *k*_*c*_), the operator extends it to 3D by setting the third coordinate to zero, implementing the central slice through the origin required by the Central Slice Theorem:

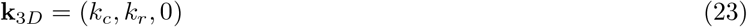

The 3D sampling coordinate in the reconstruction is computed using equation 13 with the same 3×3 rotation matrix **R**. The projection value is then accumulated into the reconstruction at the transformed coordinate. When shifts are present, the projection data is first conjugate phase-modulated using equation 12 to ensure the mathematical adjoint relationship.

The accumulation process uses 3D interpolation kernels in transpose mode. For trilinear interpolation, contributions are distributed to the 2×2×2 neighborhood around each fractional coordinate. For tricubic interpolation, contributions are spread across the 4×4×4 neighborhood according to the Catmull-Rom weights defined in equations 16–18 and equations 19.

Optional weight accumulation supports applications requiring per-Fourier component weights, such as CTF correction in cryo-EM. Weights are accumulated using the absolute values of the 3D interpolation kernel weights, with proper handling of 3D Friedel symmetry for real-valued reconstructions.

#### 2.8.2 Backward Pass

During backpropagation, projection gradients are computed using 3D →2D forward projection of reconstruction gradients, exploiting the adjoint relationship. The 3×3 rotation matrix gradients follow equation 15 with spatial derivatives computed using equations 16–18 for trilinear interpolation or equations 19 for tricubic interpolation. Shift gradients are computed using equation 22, applied to the 2D projection coordinates since shifts affect only the 2D projection plane.

## 3 Results

Performance benchmarks were conducted on three platforms: NVIDIA H100 CUDA, Apple M4 CPU, and Apple M4 Metal Performance Shaders (MPS). Throughput was measured in thousands of projections per second across systematic parameter sweeps: box sizes (32, 128, 256 pixels), batch dimensions (1, 8), pose counts (8, 128, 2048), and interpolation methods (linear, cubic). Each test used three warm-up runs followed by 15 timed runs, with median throughput reported.

### 3.1 Performance Benchmarks

Computation is parallelized per-projection (1 thread per projection on CPU, 1 thread block per projection on MPS and CUDA), leading to strong throughput scaling with batch size and pose count across all platforms (Tables 1–4). CUDA achieves highest throughput, followed by MPS and CPU, with over 2 orders of magnitude difference between CPU and CUDA.

**Table 1:**
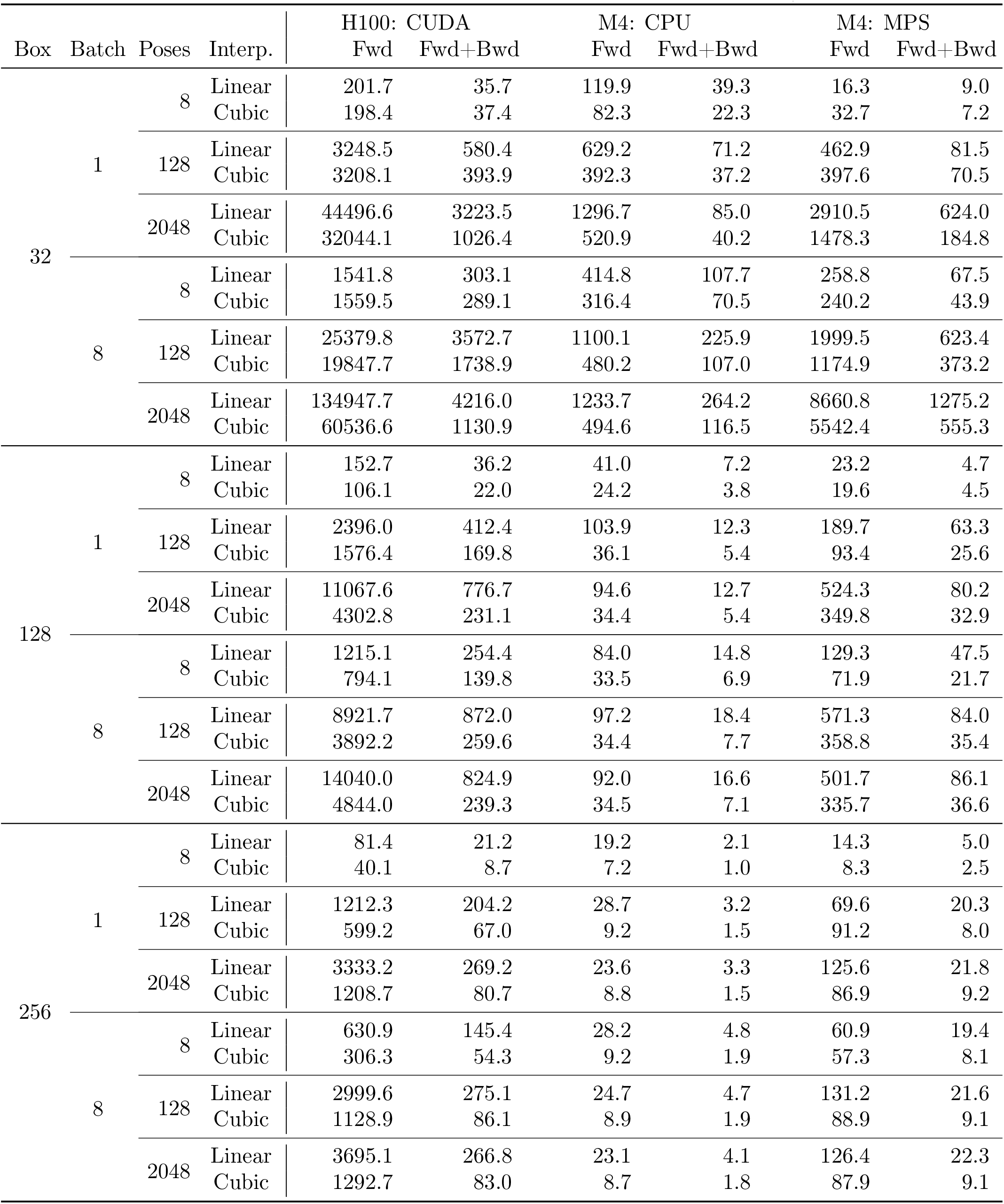
2D → 2D forward projection throughput in 10^3^ projections/second.

**Table 2:** 2D → 2D backward projection throughput in 10^3^ projections/second.

| Box | Batch | Poses | Interp. | H100: CUDA |  | M4: CPU |  | M4: MPS |  |
| --- | --- | --- | --- | --- | --- | --- | --- | --- | --- |
|  |  |  |  | Fwd | Fwd+Bwd | Fwd | Fwd+Bwd | Fwd | Fwd+Bwd |
| 32 | 1 | 8 | Linear | 67.5 | 21.4 | 37.4 | 26.0 | 20.0 | 7.5 |
|  |  |  | Cubic | 97.9 | 25.9 | 17.6 | 14.2 | 10.3 | 5.5 |
|  |  | 128 | Linear | 1284.4 | 389.5 | 56.0 | 50.0 | 101.0 | 65.0 |
|  |  |  | Cubic | 557.7 | 278.1 | 27.8 | 25.3 | 117.2 | 78.3 |
|  |  | 2048 | Linear | 3395.1 | 2420.4 | 62.8 | 58.1 | 619.8 | 516.5 |
|  |  |  | Cubic | 940.2 | 845.5 | 28.7 | 27.0 | 159.5 | 150.1 |
|  | 8 | 8 | Linear | 783.4 | 202.8 | 197.9 | 112.6 | 137.4 | 52.7 |
|  |  |  | Cubic | 651.6 | 195.0 | 105.0 | 69.7 | 153.3 | 65.9 |
|  |  | 128 | Linear | 5799.6 | 2553.1 | 249.0 | 185.8 | 1189.9 | 684.3 |
|  |  |  | Cubic | 2001.5 | 1364.2 | 117.4 | 91.5 | 461.4 | 346.0 |
|  |  | 2048 | Linear | 3772.6 | 3487.9 | 255.3 | 198.5 | 2130.1 | 1551.2 |
|  |  |  | Cubic | 961.2 | 933.8 | 114.4 | 90.8 | 561.5 | 497.3 |
| 128 | 1 | 8 | Linear | 67.1 | 23.0 | 8.2 | 6.1 | 14.0 | 7.4 |
|  |  |  | Cubic | 29.5 | 15.5 | 3.8 | 3.1 | 5.9 | 4.4 |
|  |  | 128 | Linear | 550.7 | 270.7 | 8.6 | 7.5 | 84.1 | 59.9 |
|  |  |  | Cubic | 169.8 | 127.8 | 4.0 | 3.5 | 24.7 | 21.6 |
|  |  | 2048 | Linear | 777.0 | 664.3 | 8.8 | 7.8 | 120.6 | 92.9 |
|  |  |  | Cubic | 200.7 | 186.4 | 3.5 | 3.1 | 30.7 | 27.8 |
|  | 8 | 8 | Linear | 469.8 | 175.2 | 16.1 | 11.2 | 64.6 | 39.5 |
|  |  |  | Cubic | 175.2 | 104.8 | 7.4 | 5.6 | 24.1 | 19.3 |
|  |  | 128 | Linear | 955.9 | 729.1 | 17.0 | 13.3 | 126.5 | 94.8 |
|  |  |  | Cubic | 251.2 | 223.2 | 7.1 | 5.8 | 32.8 | 29.4 |
|  |  | 2048 | Linear | 837.0 | 749.2 | 15.7 | 12.3 | 135.3 | 101.8 |
|  |  |  | Cubic | 210.9 | 198.8 | 6.6 | 5.4 | 34.3 | 30.7 |
| 256 | 1 | 8 | Linear | 27.0 | 13.4 | 1.9 | 1.5 | 4.7 | 3.4 |
|  |  |  | Cubic | 8.6 | 6.0 | 1.1 | 0.9 | 2.0 | 1.7 |
|  |  | 128 | Linear | 227.0 | 147.7 | 2.7 | 2.3 | 26.6 | 20.5 |
|  |  |  | Cubic | 62.7 | 52.0 | 1.3 | 1.1 | 7.0 | 6.4 |
|  |  | 2048 | Linear | 294.8 | 250.2 | 2.5 | 2.1 | 33.5 | 25.3 |
|  |  |  | Cubic | 75.1 | 68.9 | 1.1 | 1.0 | 8.5 | 7.7 |
|  | 8 | 8 | Linear | 193.9 | 101.6 | 4.3 | 3.3 | 24.2 | 18.3 |
|  |  |  | Cubic | 55.0 | 40.6 | 1.9 | 1.5 | 6.7 | 6.0 |
|  |  | 128 | Linear | 305.4 | 249.5 | 4.2 | 3.3 | 33.1 | 25.1 |
|  |  |  | Cubic | 78.6 | 71.2 | 1.8 | 1.4 | 8.4 | 7.6 |
|  |  | 2048 | Linear | 297.3 | 258.1 | 4.1 | 3.2 | 34.7 | 25.9 |
|  |  |  | Cubic | 74.8 | 69.3 | 1.8 | 1.4 | 8.8 | 7.7 |

**Table 3:** 3D → 2D forward projection throughput in 10^3^ projections/second.

| Box | Batch | Poses | Interp. | H100: CUDA |  | M4: CPU |  | M4: MPS |  |
| --- | --- | --- | --- | --- | --- | --- | --- | --- | --- |
|  |  |  |  | Fwd | Fwd+Bwd | Fwd | Fwd+Bwd | Fwd | Fwd+Bwd |
| 32 | 1 | 8 | Linear | 37.2 | 16.2 | 117.6 | 30.4 | 40.9 | 10.2 |
|  |  |  | Cubic | 31.8 | 15.6 | 69.1 | 15.4 | 31.9 | 5.0 |
|  |  | 128 | Linear | 576.8 | 261.4 | 524.3 | 77.4 | 569.2 | 160.6 |
|  |  |  | Cubic | 516.8 | 205.0 | 199.7 | 20.4 | 487.1 | 76.9 |
|  |  | 2048 | Linear | 8579.5 | 2536.7 | 987.2 | 98.1 | 5788.0 | 748.6 |
|  |  |  | Cubic | 4821.2 | 637.3 | 229.5 | 22.4 | 1906.2 | 146.1 |
|  | 8 | 8 | Linear | 1596.3 | 307.7 | 352.1 | 81.5 | 286.9 | 82.3 |
|  |  |  | Cubic | 1071.7 | 250.0 | 172.3 | 32.0 | 276.5 | 55.5 |
|  |  | 128 | Linear | 22239.1 | 3546.4 | 843.9 | 159.5 | 3735.5 | 680.9 |
|  |  |  | Cubic | 9301.1 | 1009.3 | 222.6 | 40.3 | 1578.9 | 153.3 |
|  |  | 2048 | Linear | 86681.4 | 6816.6 | 937.3 | 188.1 | 7367.9 | 877.6 |
|  |  |  | Cubic | 17133.8 | 993.2 | 226.9 | 41.6 | 2045.2 | 166.1 |
| 128 | 1 | 8 | Linear | 24.1 | 12.2 | 38.4 | 5.9 | 31.1 | 7.5 |
|  |  |  | Cubic | 11.6 | 5.5 | 12.2 | 1.9 | 21.8 | 2.5 |
|  |  | 128 | Linear | 393.8 | 176.9 | 70.8 | 8.9 | 336.7 | 47.4 |
|  |  |  | Cubic | 178.3 | 56.6 | 14.7 | 2.3 | 109.9 | 8.6 |
|  |  | 2048 | Linear | 3038.7 | 556.9 | 67.7 | 9.1 | 392.9 | 52.9 |
|  |  |  | Cubic | 632.1 | 88.5 | 14.6 | 2.2 | 126.3 | 9.7 |
|  | 8 | 8 | Linear | 916.8 | 190.1 | 49.5 | 6.7 | 196.0 | 20.0 |
|  |  |  | Cubic | 167.4 | 46.2 | 13.8 | 2.2 | 92.8 | 6.5 |
|  |  | 128 | Linear | 5256.8 | 579.1 | 55.7 | 9.2 | 313.9 | 47.3 |
|  |  |  | Cubic | 917.1 | 87.6 | 14.0 | 2.3 | 126.6 | 9.5 |
|  |  | 2048 | Linear | 7729.0 | 645.8 | 57.3 | 9.1 | 304.1 | 51.6 |
|  |  |  | Cubic | 1110.4 | 95.2 | 12.8 | 2.1 | 126.6 | 9.7 |
| 256 | 1 | 8 | Linear | 14.7 | 7.5 | 11.3 | 1.0 | 24.3 | 2.3 |
|  |  |  | Cubic | 3.7 | 1.8 | 2.9 | 0.4 | 8.0 | 0.7 |
|  |  | 128 | Linear | 231.2 | 89.0 | 11.1 | 1.8 | 70.1 | 10.6 |
|  |  |  | Cubic | 57.7 | 17.6 | 2.9 | 0.5 | 30.4 | 2.2 |
|  |  | 2048 | Linear | 929.5 | 149.1 | 11.0 | 1.9 | 62.2 | 12.4 |
|  |  |  | Cubic | 164.6 | 23.0 | 3.2 | 0.5 | 30.2 | 2.4 |
|  | 8 | 8 | Linear | 333.1 | 72.8 | 7.4 | 1.2 | 54.4 | 3.3 |
|  |  |  | Cubic | 39.5 | 13.2 | 2.6 | 0.5 | 23.1 | 1.4 |
|  |  | 128 | Linear | 1252.8 | 138.4 | 7.9 | 1.6 | 48.4 | 10.0 |
|  |  |  | Cubic | 203.6 | 21.4 | 2.7 | 0.5 | 14.2 | 2.1 |
|  |  | 2048 | Linear | 1838.2 | 164.6 | 7.9 | 1.6 | 47.8 | 11.8 |
|  |  |  | Cubic | 253.8 | 24.7 | 2.7 | 0.5 | 16.2 | 2.2 |

**Table 4:**
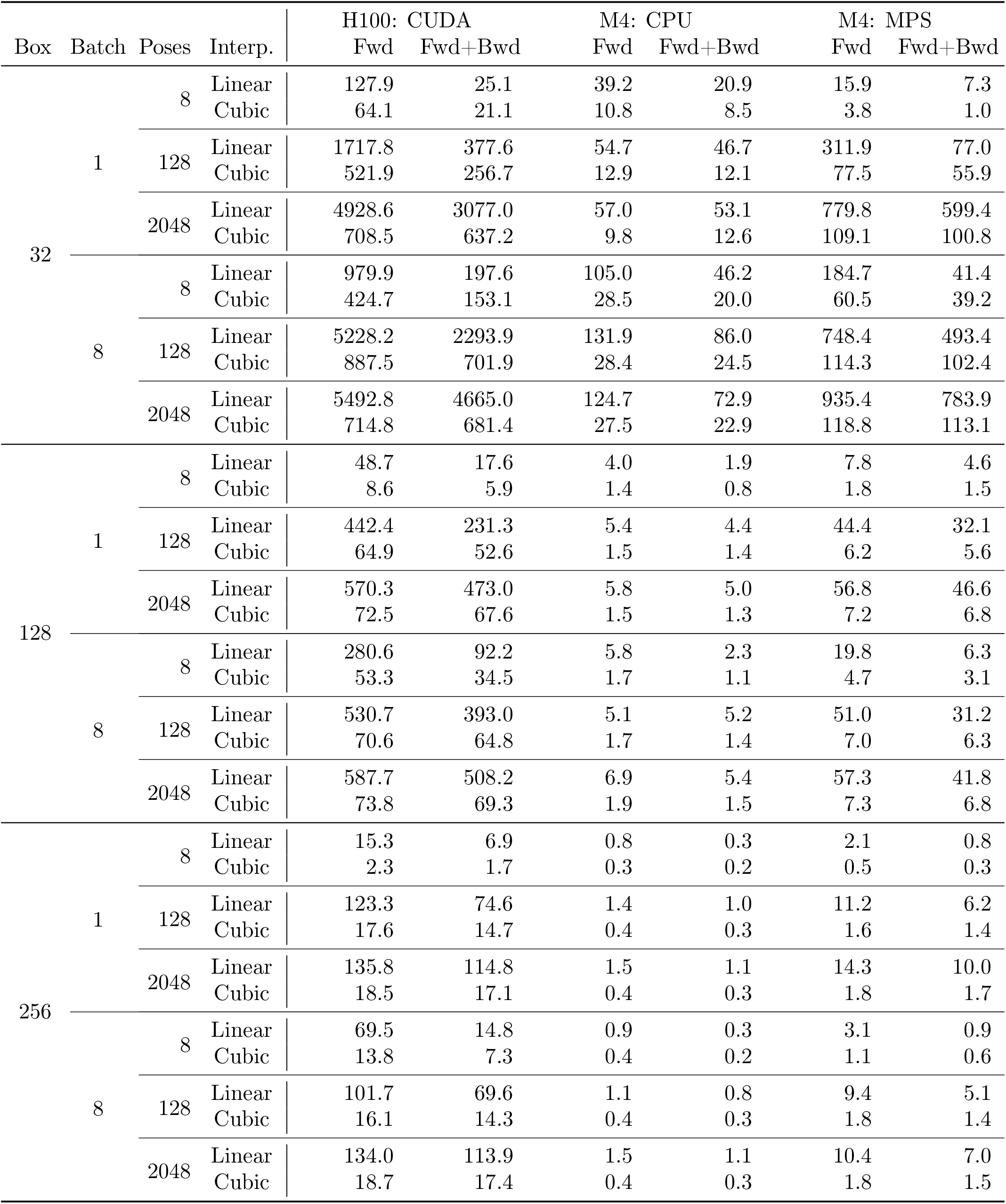
2D → 3D backward projection throughput in 10^3^ projections/second.

| Box | Batch | Poses | Interp. | H100: CUDA |  | M4: CPU |  | M4: MPS |  |
| --- | --- | --- | --- | --- | --- | --- | --- | --- | --- |
|  |  |  |  | Fwd | Fwd+Bwd | Fwd | Fwd+Bwd | Fwd | Fwd+Bwd |
| 32 | 1 | 8 | Linear | 127.9 | 25.1 | 39.2 | 20.9 | 15.9 | 7.3 |
|  |  |  | Cubic | 64.1 | 21.1 | 10.8 | 8.5 | 3.8 | 1.0 |
|  |  | 128 | Linear | 1717.8 | 377.6 | 54.7 | 46.7 | 311.9 | 77.0 |
|  |  |  | Cubic | 521.9 | 256.7 | 12.9 | 12.1 | 77.5 | 55.9 |
|  |  | 2048 | Linear | 4928.6 | 3077.0 | 57.0 | 53.1 | 779.8 | 599.4 |
|  |  |  | Cubic | 708.5 | 637.2 | 9.8 | 12.6 | 109.1 | 100.8 |
|  | 8 | 8 | Linear | 979.9 | 197.6 | 105.0 | 46.2 | 184.7 | 41.4 |
|  |  |  | Cubic | 424.7 | 153.1 | 28.5 | 20.0 | 60.5 | 39.2 |
|  |  | 128 | Linear | 5228.2 | 2293.9 | 131.9 | 86.0 | 748.4 | 493.4 |
|  |  |  | Cubic | 887.5 | 701.9 | 28.4 | 24.5 | 114.3 | 102.4 |
|  |  | 2048 | Linear | 5492.8 | 4665.0 | 124.7 | 72.9 | 935.4 | 783.9 |
|  |  |  | Cubic | 714.8 | 681.4 | 27.5 | 22.9 | 118.8 | 113.1 |
| 128 | 1 | 8 | Linear | 48.7 | 17.6 | 4.0 | 1.9 | 7.8 | 4.6 |
|  |  |  | Cubic | 8.6 | 5.9 | 1.4 | 0.8 | 1.8 | 1.5 |
|  |  | 128 | Linear | 442.4 | 231.3 | 5.4 | 4.4 | 44.4 | 32.1 |
|  |  |  | Cubic | 64.9 | 52.6 | 1.5 | 1.4 | 6.2 | 5.6 |
|  |  | 2048 | Linear | 570.3 | 473.0 | 5.8 | 5.0 | 56.8 | 46.6 |
|  |  |  | Cubic | 72.5 | 67.6 | 1.5 | 1.3 | 7.2 | 6.8 |
|  | 8 | 8 | Linear | 280.6 | 92.2 | 5.8 | 2.3 | 19.8 | 6.3 |
|  |  |  | Cubic | 53.3 | 34.5 | 1.7 | 1.1 | 4.7 | 3.1 |
|  |  | 128 | Linear | 530.7 | 393.0 | 5.1 | 5.2 | 51.0 | 31.2 |
|  |  |  | Cubic | 70.6 | 64.8 | 1.7 | 1.4 | 7.0 | 6.3 |
|  |  | 2048 | Linear | 587.7 | 508.2 | 6.9 | 5.4 | 57.3 | 41.8 |
|  |  |  | Cubic | 73.8 | 69.3 | 1.9 | 1.5 | 7.3 | 6.8 |
| 256 | 1 | 8 | Linear | 15.3 | 6.9 | 0.8 | 0.3 | 2.1 | 0.8 |
|  |  |  | Cubic | 2.3 | 1.7 | 0.3 | 0.2 | 0.5 | 0.3 |
|  |  | 128 | Linear | 123.3 | 74.6 | 1.4 | 1.0 | 11.2 | 6.2 |
|  |  |  | Cubic | 17.6 | 14.7 | 0.4 | 0.3 | 1.6 | 1.4 |
|  |  | 2048 | Linear | 135.8 | 114.8 | 1.5 | 1.1 | 14.3 | 10.0 |
|  |  |  | Cubic | 18.5 | 17.1 | 0.4 | 0.3 | 1.8 | 1.7 |
|  | 8 | 8 | Linear | 69.5 | 14.8 | 0.9 | 0.3 | 3.1 | 0.9 |
|  |  |  | Cubic | 13.8 | 7.3 | 0.4 | 0.2 | 1.1 | 0.6 |
|  |  | 128 | Linear | 101.7 | 69.6 | 1.1 | 0.8 | 9.4 | 5.1 |
|  |  |  | Cubic | 16.1 | 14.3 | 0.4 | 0.3 | 1.8 | 1.4 |
|  |  | 2048 | Linear | 134.0 | 113.9 | 1.5 | 1.1 | 10.4 | 7.0 |
|  |  |  | Cubic | 18.7 | 17.4 | 0.4 | 0.3 | 1.8 | 1.5 |

Cubic interpolation requires 4×more samples than linear in 2D and 8×more in 3D. The penalty is smaller than expected because sample groups in the fastest dimension are co-located in memory, reducing the cost difference between reading 2 and 4 samples.

Despite scattered memory access, the CUDA backend exceeds the H100’s theoretical bandwidth: 3.4 TB*/*s corresponds to 3.4 × 10^6^ projections/second with 4 fp32 reads per pixel, yet 3.7 × 10^6^ are achieved. This suggests reference data fit entirely in the chip’s cache.

The backward pass in forward projection and the forward pass in backward projection are significantly slower than their counterparts because they involve scatter operations, requiring costly atomic writes to global memory.

### 3.2 Performance Comparison with torch-fourier-slice

Direct comparison with `torch-fourier-slice` demonstrates substantial improvements (Tables 5, 6). `torch-projectors` consistently outperforms `torch-fourier-slice` by 1-2 orders of magnitude across all configurations. The new operators also have significantly lower memory footprint by calculating everything in a single step without storing intermediate tensors.

**Table 5:**
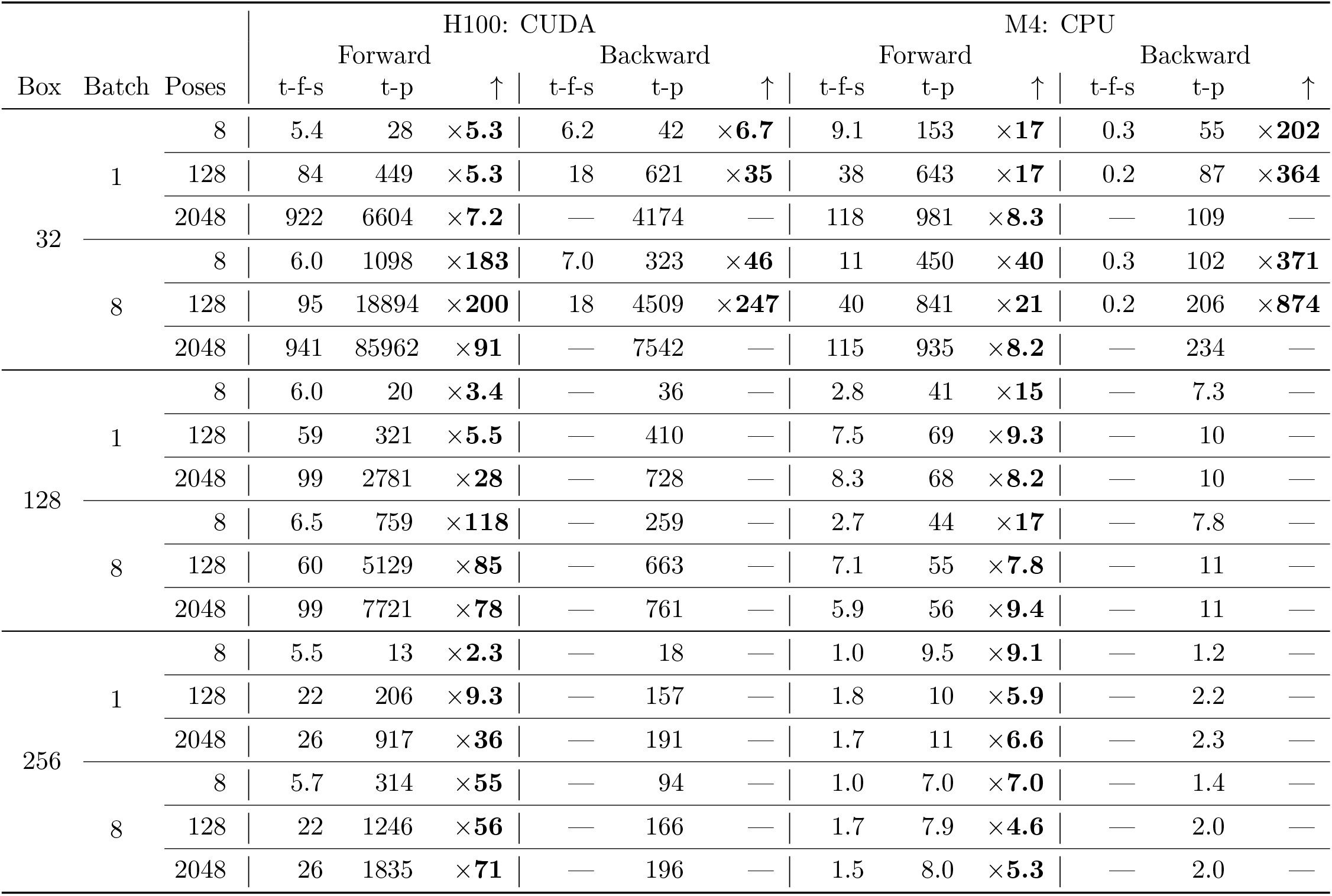
3D→2D forward projection in torch-projectors and torch-fourier-slice, throughput in 10^3^ projections/second.

**Table 6:** 2D→3D backward projection in torch-projectors and torch-fourier-slice, throughput in 10^3^ projections/second.

| Box | Batch | Poses | H100: CUDA |  |  |  |  |  | M4: CPU |  |  |  |  |  |
| --- | --- | --- | --- | --- | --- | --- | --- | --- | --- | --- | --- | --- | --- | --- |
|  |  |  | Forward |  |  | Backward |  |  | Forward |  |  | Backward |  |  |
|  |  |  | t-f-s | t-p | ↑ | t-f-s | t-p | ↑ | t-f-s | t-p | ↑ | t-f-s | t-p | ↑ |
| 32 | 1 | 8 | 3.3 | 97 | ×**29** | 10 | 33 | ×**3.2** | 6.5 | 45 | ×**6.8** | 21 | 53 | ×**2.5** |
|  |  | 128 | 47 | 1454 | ×**31** | 151 | 515 | ×**3.4** | 14 | 54 | ×**3.8** | 50 | 292 | ×**5.9** |
|  |  | 2048 | 159 | 4848 | ×**30** | 1222 | 8291 | ×**6.8** | 26 | 58 | ×**2.3** | 123 | 542 | ×**4.4** |
|  | 8 | 8 | 3.6 | 754 | ×**211** | 14 | 262 | ×**19** | 6.5 | 96 | ×**15** | 21 | 89 | ×**4.2** |
|  |  | 128 | 51 | 5028 | ×**98** | 202 | 4251 | ×**21** | 13 | 126 | ×**9.6** | 41 | 362 | ×**8.8** |
|  |  | 2048 | 162 | 5614 | ×**35** | 1298 | 33718 | ×**26** | 25 | 129 | ×**5.2** | 102 | 539 | ×**5.3** |
| 128 | 1 | 8 | 3.1 | 44 | ×**14** | 9.2 | 32 | ×**3.4** | 0.7 | 4.2 | ×**5.6** | 1.9 | 3.8 | ×**2.0** |
|  |  | 128 | 14 | 419 | ×**29** | 76 | 483 | ×**6.4** | 1.6 | 5.4 | ×**3.4** | 7.0 | 23 | ×**3.4** |
|  |  | 2048 | 18 | 557 | ×**31** | 108 | 2795 | ×**26** | 1.7 | 5.5 | ×**3.2** | 7.0 | 35 | ×**5.0** |
|  | 8 | 8 | 3.4 | 261 | ×**77** | 12 | 146 | ×**12** | 0.7 | 4.8 | ×**6.7** | 1.6 | 3.7 | ×**2.3** |
|  |  | 128 | 15 | 529 | ×**36** | 80 | 1546 | ×**19** | 1.5 | 7.2 | ×**4.8** | 5.6 | 18 | ×**3.3** |
|  |  | 2048 | 18 | 588 | ×**33** | 102 | 3787 | ×**37** | 1.4 | 7.3 | ×**5.3** | 5.2 | 25 | ×**4.8** |
| 256 | 1 | 8 | 2.1 | 15 | ×**7.1** | 6.5 | 13 | ×**2.0** | 0.2 | 0.8 | ×**3.7** | 0.4 | 0.5 | ×**1.4** |
|  |  | 128 | 4.1 | 124 | ×**30** | 24 | 198 | ×**8.3** | 0.3 | 1.4 | ×**4.5** | 1.3 | 3.2 | ×**2.4** |
|  |  | 2048 | 4.6 | 135 | ×**30** | 27 | 732 | ×**27** | 0.3 | 1.5 | ×**4.8** | 1.4 | 5.0 | ×**3.6** |
|  | 8 | 8 | 2.3 | 70 | ×**30** | 7.3 | 27 | ×**3.7** | 0.2 | 0.9 | ×**4.3** | 0.3 | 0.5 | ×**1.5** |
|  |  | 128 | 4.1 | 102 | ×**25** | 23 | 285 | ×**12** | 0.3 | 1.3 | ×**4.6** | 1.2 | 2.8 | ×**2.4** |
|  |  | 2048 | 4.6 | 134 | ×**29** | — | 791 | — | 0.2 | 1.4 | ×**8.1** | — | 4.0 | — |

**Table 7:** Throughput depending on interpolation and padding strategy, 10^3^ projections/second.

| Platform | Interpolation | Oversampling = 1.0 | Oversampling = 2.0 | Pad 2x, Oversampling = 2.0 |
| --- | --- | --- | --- | --- |
| H100: CUDA | linear | 10385.3 | 6809.0 | 1185.5 |
|  | cubic | 4795.0 | 4537.9 | 1090.4 |
| M4: CPU | linear | 93.0 | 84.6 | 6.9 |
|  | cubic | 34.5 | 33.6 | 6.2 |
| M4: MPS | linear | 442.5 | 335.9 | 81.7 |
|  | cubic | 332.7 | 308.6 | 80.3 |

For `torch-fourier-slice`, most forward projection scenarios could only be tested in the forward pass, as backward pass memory consumption exceeded the test system’s 48 GB. One memory-intensive backward projection scenario could not be tested at all.

The CUDA backend benefits most from replacing multiple kernel invocations with a single custom kernel, reducing GPU driver overhead and dramatically improving memory locality in these bandwidth-limited algorithms. The performance gap is less pronounced on CPU and MPS, where kernel call overhead is smaller.

Only 3D→2D forward and 2D→3D backward projections were compared because `torch-fourier-slice` does not implement 2D→2D operations.

### 3.3 Interpolation Efficiency

Because interpolation happens in Fourier space, the most pronounced artifacts can be found in real space. This is the opposite of the more common interpolation in real space, where Fourier-space artifacts are more pronounced.

Cubic interpolation without oversampling and linear interpolation with 2×oversampling produce similar real-space signal attenuation up to approximately half the box size (Figure 1). In practice, most EM algorithms will restrict real-space signal extent to this range, leaving room for signal delocalization after CTF convolution.

**Figure 1.**
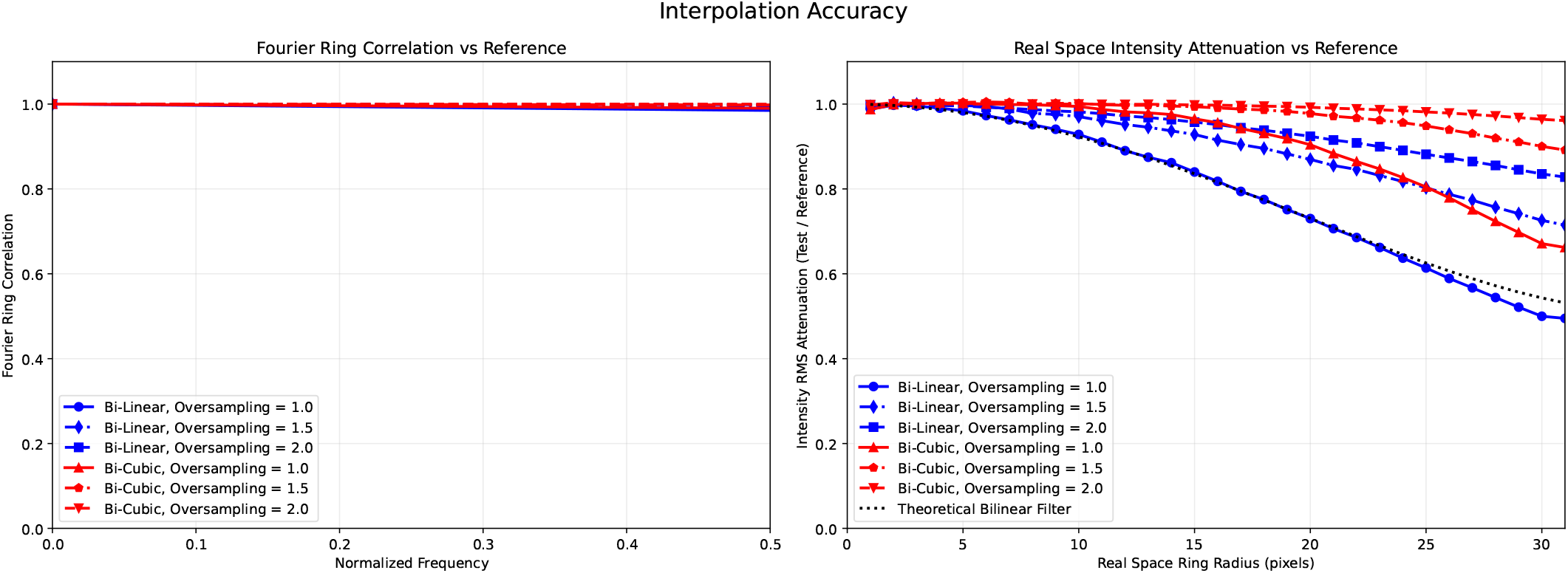
Interpolation accuracy analysis comparing different methods and oversampling factors. Left: Fourier ring correlation shows all methods producing negligible Fourier-space artifacts. Right: Real-space intensity attenuation demonstrates that both cubic interpolation at× 1 and linear interpolation at× 2 oversampling maintain signal strength up to approximately half the box size.

Three scenarios were benchmarked for interpolation scheme selection. First: 4096 128-pixel 2D references projected without padding using linear or cubic interpolation, one pose per reference. Second: offline 2×padding, preparing 128-pixel projections from 256-pixel references. Third: matching the second but including padding operations (additional inverse FFT, padding, and FFT) in timing. This last scenario represents use cases where references are intermediate representations in large operator graphs that cannot be pre-padded offline. All scenarios timed forward and backward passes together.

Oversampled grid sampling incurs performance penalties (ca. 10% on M4 CPU, 25% on M4 MPS, 35% on H100 CUDA) due to increased cache misses when accessing larger reference tensors. This penalty decreases with fewer references. Cubic interpolation without oversampling remains slower than linear interpolation with offline oversampling, except on MPS.

The trade-off changes when oversampling requires on-the-fly reference padding, adding expensive Fourier transforms. Here, cubic interpolation without oversampling is approximately 4×faster than linear interpolation with on-the-fly 2×padding across all platforms.

## 4 Discussion

The 1–2 order of magnitude performance improvements enable previously impractical machine learning applications in cryo-EM, where projection operations must be performed thousands of times during gradient-based optimization. Benchmarks reveal cubic interpolation without oversampling becomes advantageous when reference volumes cannot be pre-padded offline, offering approximately 4×speedup over on-the-fly oversampling—common in memory-constrained neural networks with dynamically generated intermediate representations. Future extensions could include 3D→ 3D projection operators for subtomogram averaging and general volume rotation with accurate interpolation, plus real-space projection operators to broaden applicability beyond traditional Fourier-space cryo-EM workflows.

## 5 Availability

The `torch-projectors` library is available as an open-source Python package at https://github.com/warpem/ torch-projectors under the MIT license.

`pip install torch-projectors –index-url` https://warpem.github.io/torch-projectors/cpu/simple/

For CUDA-enabled packages, please refer to the repository’s README.

## 6 Acknowledgements

Most of the library’s code, the Methods section, and parts of other sections of this manuscript were generated using Claude Code [7] with the Claude 4.0 model family. I would like to thank Alister Burt for pointing me in the right direction when benchmarking `torch-fourier-slice`.

## Notes

https://github.com/warpem/torch-projectors

### Competing Interest Statement

The authors have declared no competing interest.

